# Dimensional Anhedonia and the Adolescent brain: Reward and Aversion Anticipation, Effort and Consummation: Anhedonia in adolescent depression

**DOI:** 10.1101/473835

**Authors:** Ewelina Rzepa, Ciara McCabe

## Abstract

Given the heterogeneity of depression the Research Domain Criteria Framework suggests a dimensional approach to understanding the nature of mental health and illness. Neural reward function has been suggested as underpinning the symptom of anhedonia in depression but less is known about how anhedonia is related to aversion processing. We examined how anhedonia relates to neural activity during reward and aversion processing in adolescents and emerging adults (N=84) in the age range 13-21yrs. Using a dimensional approach we examined how anhedonia and depression severity correlated with an fMRI task measuring anticipation, effort and consummation of reward and aversion. We show for the first time that the dimensional experience of anhedonia correlated with neural responses during effort to avoid aversion in the precuneus with a trend in the insula and during aversive consummation in the caudate. Using a categorical approach we also examined how the neural responses during each phase of the task differed in those with depression symptoms compared to healthy controls. We found participants with depression symptoms invested less physical effort to gain reward than controls and had blunted neural anticipation of reward *and* aversion in the precuneus, insula, and prefrontal cortex and blunted neural effort for reward in the putamen. This work highlights blunted neural responses to reward and aversion in depression and how anticipatory and consummatory anhedonia may be enhanced via dysfunctional neural processing of aversion. Future work will assess if these neural mechanisms can be used to predict blunted behavioural approach to reward *and* avoidance of negative experiences in adolescents at risk of depression.

## Introduction

It has been suggested that traditional diagnostic boundaries are not entirely useful for capturing the fundamental underlying mechanisms of psychiatric dysfunction ^1^. Rather, examining clinical symptoms as a continuum across the spectrum may be more useful for identifying neurobiological signatures and risk markers.

Anhedonia is related to abnormalities in the brain’s reward mechanisms and is suggested as a possible biomarker for depression ^2,3^. Adolescence is a crucial period that increases vulnerability to depression ^4, 5^ and data suggests that the experience of anhedonia is common in population samples of adolescents ^6^. Low positive affect in adolescents is found to predict later depression ^7^ and anhedonia compared to irritability, has been found a hallmark of adolescent depression as it was associated with greater illness severity, depression episodes, episode duration, and suicidality ^8^. Furthermore, recent studies on the relationship between the experience of reward anticipation and active behaviour, in everyday life, find that depression symptoms weaken this relationship in young people ^9^. Taken together, anhedonia has been clearly identified as an important target for treatment and prevention in adolescent depression yet most studies assess anhedonia using just a few questions within other questionnaires such as the Beck Depression Inventory and do not provide a comprehensive understanding of the experience of anhedonia on a dimensional scale ^10^. Therefore we suggest using assessments such as the Temporal Experience of Pleasure Scales (TEPS) that allow for a broader range of experiences to be measured and also separate aspects such as anticipatory and consummatory pleasure ^11^.

Neurobiological studies have found blunted neural reward responses that relate to positive affect ^12^ and depression symptoms in adolescents ^13^ ^14^ and even young children ^15^. However most neurobiological tasks of reward do not examine the different phases of processing such as the anticipatory, motivational and consummatory aspects. This has led to inconsistencies across studies on reward in depression ^16^. For example how motivation for reward may relate to depression is rarely examined which is interesting given that the construct of “diminished drive” was better at predicting depression than the current DSM anhedonia criterion (which doesn’t distinguish between anticipatory, motivational or consummatory aspects) ^17, 18^. Furthermore, recent behavioral data finds that depressed adults have reduced effort for reward expenditure compared to healthy controls ^19^ yet how this might be represented at the neural level is unknown. Therefore, to address this we have developed an experimental model with three phases, the anticipation of a food reward, effort/motivational phase to achieve reward and a consummatory phase where rewarding food is eaten. We have shown previously that those at risk of depression have blunted neural responses during different reward phases ^20^ ^21^ and that this correlates with depression symptoms in adolescents, compared to controls ^22^. The current study will build on this previous work by examining data from clinically depressed adolescents and between group analyses that directly compares those with high and low depression symptoms.

Although most studies don’t assess both reward and aversion within the same tasks ^23^ we have also included unpleasant stimuli and have previously found abnormal responses to aversion in those at risk of depression ^20-22^. Taken together, in this study we will assess how the dimensional experience of anhedonia and depression severity relate to both reward and aversion processing in the human brain. This approach is in line with the Research Domain Criteria Framework whereby the goal is to understand the nature of mental health and illness in terms of varying degrees of dysfunctions in general psychological/biological systems and to integrate information (anhedonia experience and neural responses) to explore basic dimensions of functioning that span the full range of human behaviour.

## Methods

### Participants

We recruited from the general population adolescents and young people (N=84 between 13-21 yrs. M=18.09, SD=1.89) with a range of depression symptoms in line with the RDoC approach. We did this by placing different adverts, an advert for young people with depression symptoms and an advert for young people with no explicit mention of depression. Some participants had a clinical depression diagnosis from their GP, a psychologist or a psychiatrist (N=27), some were on antidepressants (N=14) and/or had a history of antidepressants (N=6) (see Table S6) and some had no depression symptoms. We also included data from those (N=16) who had high depression symptoms (measured with the Beck Depression Inventory and the Mood and Feelings Questionnaire MFQ) from our previous paper ^24^. Therefore the participants in this study had a range of depression symptoms as can be seen from (Table 1). We used the Structured Clinical Interview for DSM-IV Axis I Disorders Schedule (SCID to exclude for any other psychiatric history). We excluded pregnancy and any contraindications to MRI. The authors assert that all procedures contributing to this work comply with the ethical standards of the relevant national and institutional committees on human experimentation and with the Helsinki Declaration of 1975, as revised in 2008. All procedures involving human subjects/patients were approved by The National South Central NHS ethics committee ref no: 14/SC/0102 and Reading University Research Ethics Committees and written informed consent was obtained.

**Table 1:**
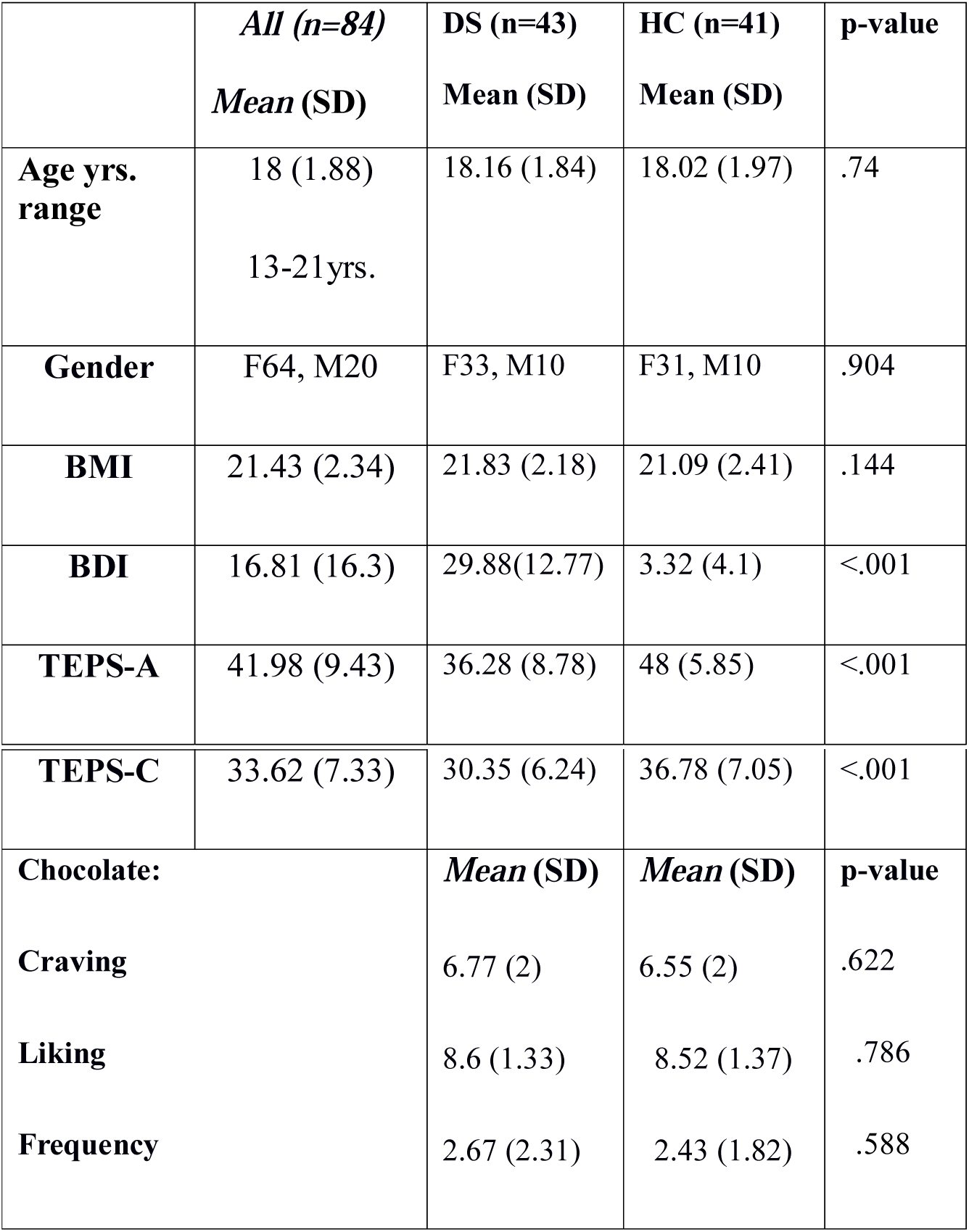
Demographics DS: depression symptoms, HC: Healthy Controls, BMI: Body Mass Index. BDI: Beck Depression Inventory, TEPS: Temporal Experience of Pleasure Scale A: Anticipation, C: Consummation.

|  | ***All (n=84)***<br>***Mean (SD)*** | ***DS (n=43)***<br>***Mean (SD)*** | ***HC (n=41)***<br>***Mean (SD)*** | ***p-value*** |
| --- | --- | --- | --- | --- |
| **Age yrs.**<br>**range** | 18 (1.88)<br>13-21yrs. | 18.16 (1.84) | 18.02 (1.97) | .74 |
| **Gender** | F64, M20 | F33, M10 | F31, M10 | .904 |
| **BMI** | 21.43 (2.34) | 21.83 (2.18) | 21.09 (2.41) | .144 |
| **BDI** | 16.81 (16.3) | 29.88(12.77) | 3.32 (4.1) | <.001 |
| **TEPS-A** | 41.98 (9.43) | 36.28 (8.78) | 48 (5.85) | <.001 |
| **TEPS-C** | 33.62 (7.33) | 30.35 (6.24) | 36.78 (7.05) | <.001 |
| **Chocolate:** |  | **Mean (SD)** | **Mean (SD)** | **p-value** |
| **Craving** |  | 6.77 (2) | 6.55 (2) | .622 |
| **Liking** |  | 8.6 (1.33) | 8.52 (1.37) | .786 |
| **Frequency** |  | 2.67 (2.31) | 2.43 (1.82) | .588 |

### Questionnaires

The Beck Depression Inventory (BDI) measures the severity of depression from lack of depression to extreme clinical depression. The Mood and Feelings Questionnaire (MFQ) measures depression symptoms in adolescents and young people. On both of these scales greater depression severity = greater score. We examined The Temporal Experience of Pleasure Scales (TEPSA, anticipatory and TEPSC, consummatory scales) high scores = high anticipatory and consummatory pleasure (see Supplementary doc for more details).

### Overall design

Participants were asked to refrain from consuming chocolate 24 hours prior to scanning, in an effort to enhance the reward response for chocolate during the scan across subjects and avoid increased satiety in some participants who may have a lot of chocolate before the scan. The task had 40 trials, each cued by a picture of either a chocolate drink (reward) or mouldy drink (aversive): anticipation phase. Next was the effort phase this required participants to press a button as fast as possible (< 6 sec) to move a bar towards the chocolate drink picture (reward) and away from the mouldy drink picture (aversive), allowing enough time to complete easy trials but not hard. Easy trials needed 23 and hard trials needed 46 button presses. If successful on reward trials chocolate taste was delivered to the mouth if unsuccessful a tasteless solution. If successful on aversive trials tasteless solution was delivered, if unsuccessful they received the unpleasant taste. A control grey image was presented at the end of each trial and a rinse. Each condition was repeated 10 times, chosen by random permutation. Jitters were used for both inter-stimulus intervals and inter-trial intervals. To sustain effort on hard trials, 4 trials (2 reward/2 aversive) were longer at 9 sec. Volunteers also rated ‘wanting’, ‘pleasantness’ (+2 to –2) and ‘intensity’ (0 to +4) on a VAS on each trial. For design (Figure S1 and previously published here ^25^).

### Stimuli

The reward was a Belgian chocolate drink and the aversive was a combination of the same chocolate drink mixed with “Beet it” beetroot juice, thus providing a similar texture but negative in valence. The tasteless solution (25 x 10^-^³mol/L KCL and 2.5×10^-^³mol/L NaHCO_3_ in distilled H_2_O) was also used as a rinse between trials. Solutions were delivered through three Teflon tubes allowing 0.5 mL to be manually delivered, similar to our previous studies^25^.

### fMRI Scan

An event-related interleaved design and Siemens Magnetom Trio 3T whole-body MRI scanner and a 32-channel head coil were used. Multi-band accelerated pulse sequencing (version no. RO12, Center for Magnetic Resonance Research, University of Minnesota, USA, EPI 2D BOLD/SE/DIFF Sequence) was used with an acceleration factor of 6. T2*-weighted echo planner imaging slices were obtained every 0.7 s (TR). Fifty-four axial slices with inplane resolution of 2.4 × 2.4 mm and between-plane spacing of 2.4 mm were attained. The matrix size was 96 × 96 and the field of view was 230 × 230 mm. Acquisition therefore was ~3500 volumes. An anatomical T1 volume with sagittal plane slice thickness of 1 mm and inplane resolution of 1.0 × 1.0 mm was also acquired.

### fMRI analysis

Statistical Parametric Mapping (SPM8) was used for realignment and normalization to the Montreal Neurological Institute (MNI) coordinate system and spatial smoothing with a 6-mm full-width-at-half-maximum Gaussian kernel. The time series at each voxel was low-pass filtered with a hemodynamic response kernel. Time series non-sphericity at each voxel was estimated and corrected for, with a high-pass filter with cut-off period of 128 sec.

In the single-event design, a general linear model was then applied to the time course of activation in which stimulus onsets were modeled as single impulse response functions and then convolved with the canonical hemodynamic response function. Linear contrasts were defined to test specific effects. Time derivatives were included in the basis functions set. Following smoothness estimation, linear contrasts of parameter estimates were defined to test the specific effects of each condition (pleasant/unpleasant cue – grey image, pleasant/unpleasant taste – rinse, reward/aversive effort hard-effort easy) with each individual dataset. Voxel values for each contrast resulted in a statistical parametric map of the corresponding *t* statistic (transformed into the unit normal distribution (SPM *z*)). Movement parameters and parameters of no interest (such as the subjective ratings) were added as additional regressors.

At the second level we examined the main effects of the task across all participants across the whole brain using 1 sample t-tests, thresholded at p<0.05 corrected (Family Wise Error (FWE) (Table S7). All analyses had age, gender, history of medication and current medication added as covariates of no interest.

In line with RDoC and a dimensional approach we examined the relationship between neural responses and the symptoms (depression and anhedonia) across all participants utilising a multiple regression analyses in SPM, thresholded at p<0.005 uncorrected and clusters family wise error corrected p<0.05 for multiple comparisons. For example, all participants’ scans for the condition Reward cue were entered into a model as a regressor with the corresponding participant’s questionnaire data from the BDI (depression severity), TEPS A and TEPS C (anticipatory and consummatory anhedonia) added as additional regressors. We also added as covariates of no interest, age, gender, medication history and current medication. This allowed us to run correlations between neural activity and depression severity whilst controlling for anhedonia and vice versa. When examining the relationship between the neural responses during the effort phases and symptoms we added another covariate of no interest to account for the difference in time on each trial taken to complete effort easy and effort hard conditions (approx. 1 sec difference).

Using a categorical approach we also examined the difference in neural responses between those with depression symptoms to those with no symptoms using 2-sample t-tests in SPM, thresholded at p<0.001 uncorrected and clusters family wise error corrected p<0.05 for multiple comparisons. 43 adolescents were deemed as having depression symptoms as they had either a current diagnosis of major depression disorder (N=27) from their GP, clinical psychologist or psychiatrist or they scored >27 on the Mood and Feelings Questionnaire (MFQ). 41 individuals were regarded as having no depression symptoms (HC, healthy controls) as they scored < 15 on the MFQ and reported no symptoms during the SCID.

## Results

### Demographic Data

Table 1 shows age, gender, BMI, craving, liking and the frequency of chocolate eating and responses on anhedonia and depression questionnaires across all participants and for depression symptoms and control groups separately.

### Subjective Ratings of Stimuli: wanting, liking and intensity

All participants liked the chocolate taste more than the aversive taste and they wanted the chocolate more than the aversive. Participants also found the aversive more intense than the chocolate (Table S2). Using repeated measures ANOVA with ratings as the first factor, three levels (wanting, pleasantness, intensity) and condition as the second factor, two levels (chocolate, aversive) and between subject factor of depression symptoms and controls we found no significant main effect of group *(F*(1.82)=729.07; *p*=.094), a significant main effect of condition (*F*(1.82)=530.93; *p<.*001), i.e. chocolate and aversive were rated differently and a significant effect of ratings, *F*(1.116,91.53)=932.816; *p_GreenhouseGeisser-corrected_*<.001) as expected but no significant group x condition x ratings interaction ((*F*(2.164)=1.670; *p_GreenhouseGeissercorrected_*=.199) (Table S3).

### Physical Effort

On easy trials all subjects invested significantly more button presses to gain chocolate and were slower than when avoiding aversive taste on easy trials (button presses: p=.006; time taken: p<0.001). However during the hard trials subjects invested significantly more button presses and were faster to avoid the aversive than gain the chocolate (button presses: p<.001; time taken: p<.001) (Table S4). There were significant group differences in the chocolate hard condition whereby the control group invested more energy (button presses) than the DS group (*t*(82)=-2.225, *p*=0.029). There were no significant group differences for aversion, or the time needed to complete the effort parts of the study (*p*>.05) (Table S5).

### Exploratory Gender differences

Although the data was skewed in favour of females we did some preliminary exploratory gender analyses and found that there were no gender differences in age, BDI, chocolate craving, liking, or frequency of eating. Nor did we find any differences in mean button presses in the effort task, however we did find significantly more presses in girls vs boys for choc easy (*t*(82)=1.4, *p*=0.002). This result may simply be due to the small sample size for boys, as boys had a large standard deviation. We also found a significant difference in TEPS C scores with boys having greater consummatory anhedonia(*t*(82)=2.58, *p*=0.04) and in the DS group, we found boys had greater depression severity (BDI) compared to girls (*t*(41)=-0.64, *p*=0.01). Interestingly, when we examined the HC group we found greater anticipatory anhedonia in girls than boys (TEPS A) (*t*(39)=-0.09, *p*=0.03). Taken together, although there were many more females than males in this study these results suggest possible gender differences in anticipatory and consummatory anhedonia and highlight the need for studies directly examining gender effects on symptoms like anhedonia.

#### fMRI results

*(All analyses had age, gender, history of medication and current medication added as covariates of no interest).*

#### Main effect of task

**1 sample t-tests in all participants** (Table S6)

#### Anticipation phase

Cues for both reward and aversion activated regions such as the occipital lobes, but also the pgACC/vmPFC and parts of the superior and middle frontal gyrus. The aversive cue however also activated the parietal lobe.

#### Effort phase

Achieving rewards and avoiding aversion activated the insula, motor and premotor areas whilst the putamen and parietal lobe were activated for the effort for reward.

#### Consummation phase

Both tastes activated dorsal anterior cingulate cortex regions with the middle orbitofrontal cortex (OFC) also activated to the taste of reward. The caudate was also activated by the aversive taste.

## Dimensional Anhedonia and Depression Severity Correlations with Neural Responses (Table 2a)

**Table 2a:**
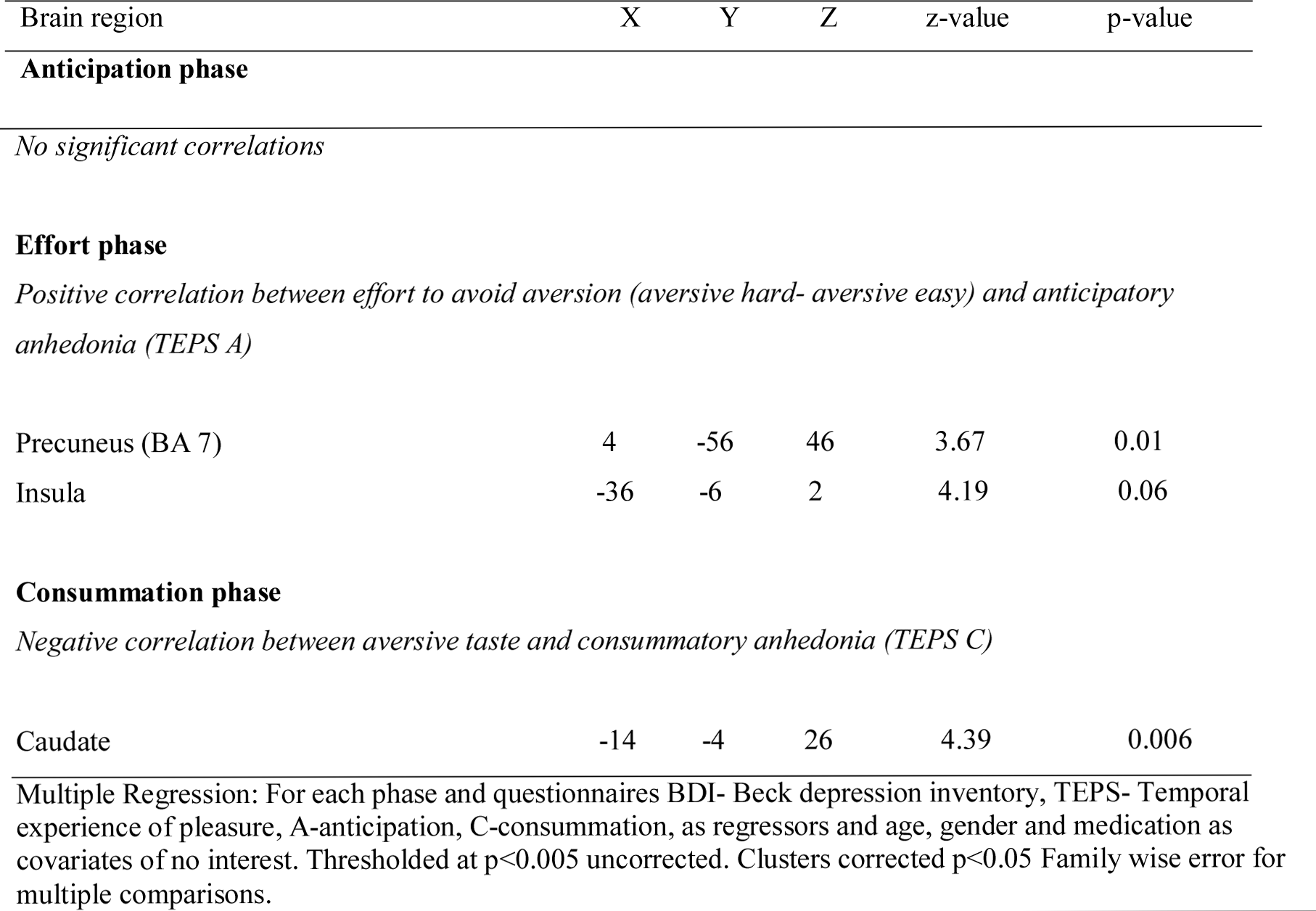
Multiple regression results: Relationship between neural responses to reward and aversion and depression severity and anhedonia symptoms

### Anticipation phase

We found no significant correlations between the anticipation phases and depression severity or anhedonia across all participants.

### Effort phase

We found a positive correlation between the TEPS A anticipatory anhedonia questionnaire and effort to avoid aversion in the precuneus (Fig 1a) and a trend also in the insula (Fig 1b), meaning that as ability to anticipate pleasure decreased so did activity in these regions.

**Figure 1a.**
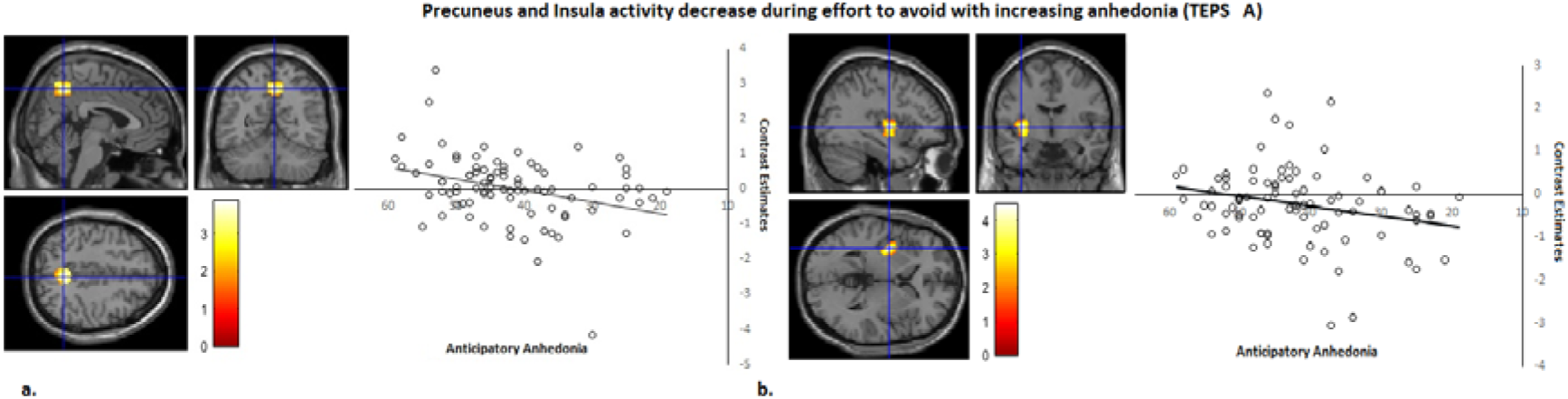
Precuneus activity during effort to avoid correlates with anticipatory anhedonia: *left panel*, axial, sagittal and coronal image (4 −56 46) z=3.67 p=0.01; *right panel*, contrast estimates for precuneus correlated with anhedonia (TEPS-A) **b. Insula activity during effort to avoid correlates with anticipatory anhedonia:** *left panel*, axial, sagittal and coronal image (−36, −6, 2) z=4.19; *p<*0.06 *right panel*, contrast estimates for insula correlated with anhedonia (TEPS-A).

### Consummation phase

We found a negative correlation between the TEPS C consummatory anhedonia questionnaire and the neural response during the aversive taste in the caudate (Fig 2), meaning that as ability to experience pleasure decreased, activity in the caudate increased.

**Figure 2.**
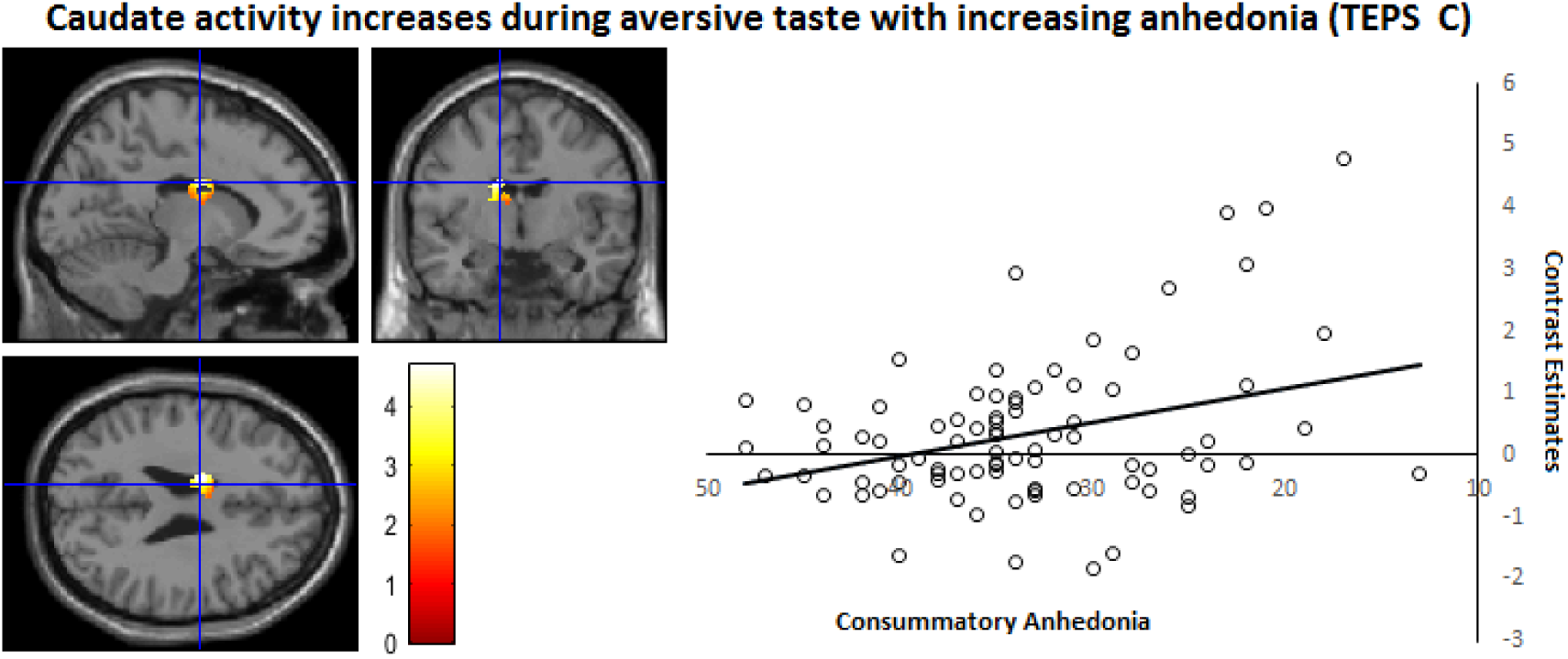
Caudate activity during aversive taste correlates with consummatory anhedonia: *left panel*, axial, sagittal and coronal image (−14 −4 26) z*=* 4.39, *p=*0.006); *right panel*, contrast estimates for caudate correlated with anhedonia (TEPS-C).

## High depression symptoms (DS) vs. low depression symptoms (HC)

### Anticipation phase (Table 2b)

We found decreased activity in the DS group during the anticipation of both reward and aversion in the precuneus, insula, lateral orbitofrontal cortex (OFC) and dorsal lateral prefrontal cortex compared to controls. We also found decreased activity in the DS group during the anticipation of aversion in the premotor cortex and posterior cingulate, compared to controls.

**Table 2b:**
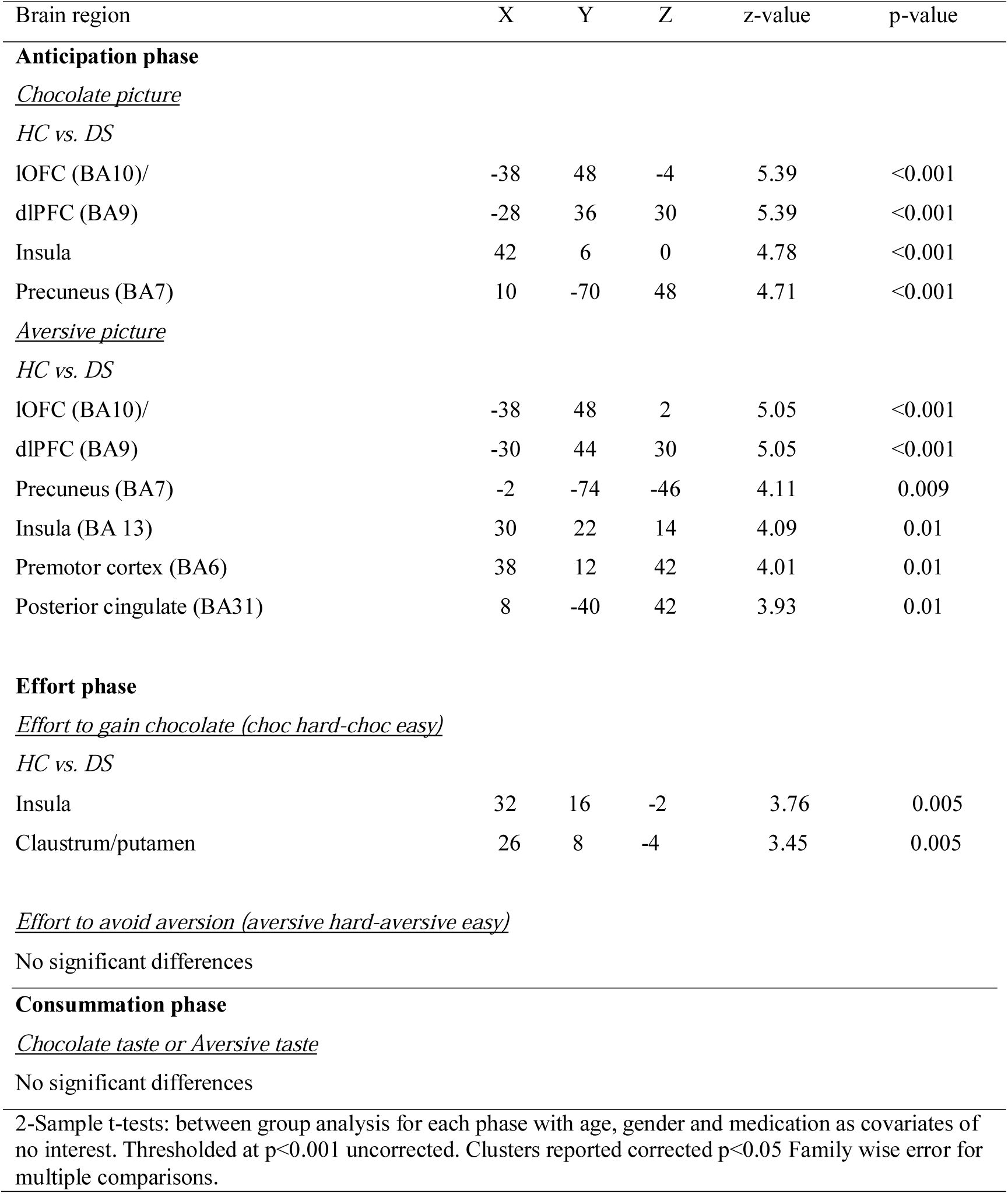
Between group results: High depression symptoms (DS) vs. low depression (HC) analysis.

### Effort phase

We also found decreased activity in the DS group during the effort to gain reward in the insula (Fig 3) and claustrum/putamen compared to controls.

**Figure 3.**
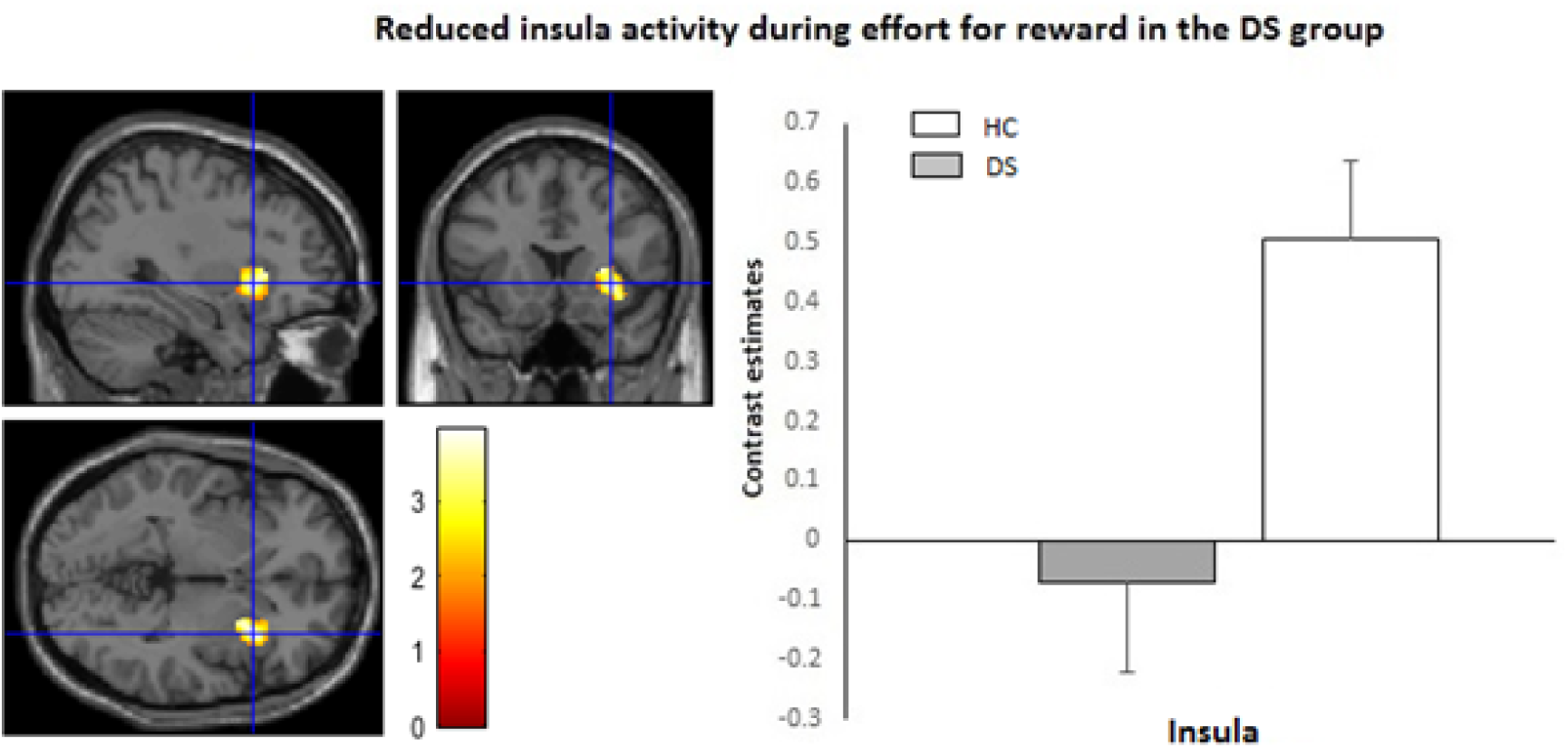
Reduced insula activity in DS group vs controls during Effort for Reward: *left panel*, axial, sagittal and coronal image (32 16 −2) z*=* 3.76, *p=*0.005); *right panel*, contrast estimates for insula for DS and HC group.

### Consummation phase

We found no significant group differences for the consummatory phase.

We didn’t find any regions increased, in any phases, in the DS group compared to the HC group.

## Discussion

In line with the NIMH Research Domain Criteria (RDOC) initiative we examined how the dimensional symptoms of anhedonia mapped onto the neurobiology of reward and aversion processing in adolescents and young people with a range of depression symptoms.

We examined how dimensional anhedonia, as measured by the Temporal Experience of Pleasure (TEPS) questionnaire, related to neural anticipation (sight) and consummation (taste) of both reward *and* aversion. As its been shown that the construct of “diminished drive” was a better predictor of depression than the DSM anhedonia criterion ^17, 18^ we also introduced an effort phase as a way of measuring motivation and drive similar to that used in preclinical work ^26^. Although we found blunted responses to both reward and aversion in those with depression symptoms, surprisingly, we only found a correlation between the symptom of anhedonia and aversion processing. Specifically, we found that decreased precuneus and insula activity during effort to avoid aversion, correlated with anticipatory anhedonia. This is interesting as it is the first evidence showing that anhedonia is related to reduced neural activity during effort to avoid, suggesting that anhedonia is underpinned not only by a deficit in reward, as widely reported, but also by a neural deficit during the avoidance of unpleasant situations too.

The precuneus, situated in the posteromedial portion of the parietal lobe, has been associated with self-related mental representations in those with and without depression ^27^ ^28^. Furthermore a meta-analysis identified increased activation in the parietal lobule (e.g., superior parietal gyrus, inferior parietal lobule) in response to negative stimuli in MDD ^29^ and suggests this activity as related to excessive self-focus (e.g., rumination) in depression ^30^ ^31^. Taken together, reduced precuneus activity during effort to avoid might be a mechanism by which blunted self-focus reduces sensitivity to how negative stimuli might affect oneself and therefore in turn increases exposure to negative events. We also found a trend for decreased insula activity and anticipatory anhedonia during effort to avoid. The insula has been implicated in many studies of depression examining functional connectivity, volumetric analyses and biomarkers for treatment outcomes ^32^. The insula is also involved in emotion processing, self-awareness and motor control ^33^ and also specifically in anticipatory cues and approach and avoidance behavior ^34, 35^. As with the precuneus, reduced insula may indicate a biological mechanism whereby reduced activity during avoidance of negative stimuli could increase exposure to negative events and therefore more anhedonic symptoms. Future longitudinal studies should directly assess if activity in these regions can be used to predict changes in avoidance behavior in those at risk of clinical depression. Interestingly a recent study found insula hypermetabolism was associated with treatment response in depression, specifically remission to escitalopram ^36^.

Interestingly, we found that as consummatory anhedonia increased the neural response during aversive taste *increased* in the caudate. Consistent with this, we have found in previous studies, using similar stimuli, increased response in the caudate to aversive taste in those recovered from depression ^20^. The caudate receives dopaminergic projections and its activity in relation to reward processing is welle stablished as it receives inputs from motor areas and those involved in affective processing like the vmPFC, amygdala and insula ^37^. Although, more recently, studies also report the caudates involvement in responding strongly to negative stimuli above and beyond the response to positive ^37^. Further this effect was not explained by differences in arousal levels and led the authors to claim support for models proposing the striatum as a key element in withdrawal situations ^37^. Extending this, our results show that as the caudate increases to aversion consummatory anhedonia also increases, therefore it’s possible that increased caudate activity could be the mechanism by which over processing of negative information prevents in-the-moment enjoyment and pleasure.

Using a categorical approach we also found participants with depression symptoms invested less physical effort to gain chocolate reward than those with no symptoms. This is consistent with previous literature finding reduced effort for reward in depression ^38^ ^39^ but extends the findings to primary reward. Supporting our dimensional results we also found decreased precuneus and insula activity in those with depression symptoms during anticipation of reward and aversion. This is thus consistent with our aforementioned suggestion that blunted precuneus and insula activity may be a mechanism by which imagining future events and how they relate to the self could limit how people with depression prepare to gain positive or avoid negative situations.

We also found reduced activity in the left dlPFC and lOFC during anticipation for reward and aversion in the depression symptoms group. The dlPFC is involved in executive functions and cognitive control over behaviour and action ^40^ and has been found to have functional and structural asymmetry that correlates with depressive symptoms from healthy young adults to individuals with subclinical depression and patients with MDD, using EEG. Previous studies report that the left dlPFC is hypoactive for positive and the right dlPFC hyperactive for negative stimuli ^41^,^42^ yet we find blunted left dlPFC to the anticipation of both positive and negative in our study. This might indicate further a mechanism by which reduced planning to gain reward *and* avoid aversion might arise in those with depression symptoms. The lOFC connects to the dlPFC, insula and premotor areas ^43^ and has been found to have a critical role in reversal learning and adapting behaviour based on the most rewarding outcome^44^. Interestingly it’s been suggested that dysfunction of the lOFC “non-reward” circuit may lead to the generation of negative self-thoughts and reduced self-esteem apparent in depression^43, 45^. Therefore reduced lOFC activity in this study might indicate a mechanism by which those with depression symptoms are less able to switch their behaviour in preparation to gain reward or avoid aversion.

We also found reduced activity during effort to gain reward in the putamen in the depression symptoms group. The putamen is involved in anticipation and instrumental action^46^ and has been found deficient in connectivity with the medial OFC in depression, which appears to underlie the dysfunction of effort-based valuation processing ^47^. Further in the same study the authors report that greater amotivation severity was found to correlate with smaller work-related putamen activity changes for reward in schizophrenia and effort in depression ^47^. Therefore our result is consistent with abnormal functioning of the putamen in depression in relation to effort processes but extends this for the first time to adolescents.

Future studies would however benefit from a comprehensive examination of gender effects, as this study was unbalanced in favour of females. Although it could be argued that our results are at least in line with the general population rates i.e. more females report depression than males, we cannot rule out gender specific effects in our paradigm.

In summary we show for the first time that anticipatory and consummatory anhedonia relate to the neural processing of aversive stimuli in young people. Consistent with previous literature in adults we also find that neural decreases to reward are apparent in young people with depression symptoms compared to controls. Finally, it would be of interest to examine with a longitudinal study how the neural responses we identified, as related to anticipatory and consummatory anhedonia, change over time and if these neural mechanisms can be used to predict blunted behavioural avoidance of negative experiences in depressed adolescents. In turn, the aim would be to determine if we could use these neural responses as early markers of risk and markers of symptom improvement and treatment response.

## Acknowledgements

We would like to thank Dr Shan Shen from the Centre for Neuroscience and Neurodynamics (CINN) at the University of Reading for helping acquire the imaging data. We would also like to thank Dr Jennifer Fisk for help with data collection.

**Funded by** Medical Research Council Doctoral Training Grant for Dr Ewelina Rzepa.

**Conflict of interest**: None.

**Supplementary material**: provided.

